# Sustained Co-evolution in a Stochastic Model of the Cancer-Immune Interaction

**DOI:** 10.1101/679746

**Authors:** Jason T. George, Herbert Levine

**Author notes:** Corresponding authors Email addresses (Jason T. George), (Herbert Levine).

## Abstract

The dynamical interaction between a growing cancer population and the adaptive immune system generates diverse evolutionary trajectories which ultimately result in tumor clearance or immune escape. Here, we create a simple mathematical model coupling T-cell recognition with an evolving cancer population which may randomly produce evasive subclones, imparting transient protection against the effector T-cells. We demonstrate that T-cell turnover declines and evasion rates together explain differential probabilities in early incidence data for almost all cancer types. Fitting the model to TRACERx evolutionary data argues in favor of substantial and sustained immune pressure exerted on a developing tumor, suggesting that measured incidence is a small proportion of all cancer initiation events. Most generally, dynamical models promise to increase our quantitative understanding of many immune escape contexts, with applications to cancer and intracellular pathogenic infections.

## 1. Introduction

T-cell immunotherapy has revolutionized modern cancer therapy, delivering durable remission outcomes with higher rates of overall survival for many cancer subtypes (Couzin-Frankel, 2013; Luke et al., 2017). The efficacy of immunotherapy stems from anti-tumor immunity imparted by the cytotoxic CD8+ T-cell repertoire, occurring as a result of effective targeting of tumor associated antigens (Coulie et al., 2014). This detection capability is consistent with theoretical expectations (George et al., 2017; Mayer et al., 2019) and has been empirically validated during natural cancer progression as well as in the post-immunotherapeutic setting (Yadav et al., 2014; Zamora et al., 2018).

Consequently, one critical phenomenon that determines ultimate tumor outcome is immunosurveillance, which relates to the degree to which tumor progression (typically in the absence of immunotherapy) is controlled by cancer-immune co-evolution. To date, a majority of our understanding of this complex interaction has been experimentally driven (Ott et al., 2017; Sahin et al., 2017; Leach et al., 1996; Sadelain et al., 2017) with the identification of immunoediting and tumor antigen negative selection during early stage disease (Fridman et al., 2011; Rosenthal et al., 2019). The resulting stochastic evolutionary trajectories are quite diverse (Dunn et al., 2002), and a mathematical framework which formalizes the underlying probabilistic dynamics governing the tumor-immune interaction promises to shed light on this complex system and may ultimately guide patient-specific immunotherapy.

The idea of modeling tumor progression and, in particular, treatment evasion is not new. Well-established stochastic models of acquired resistance have revolutionized our understanding of acquired drug resistance in the setting of targeted therapy (Iwasa et al., 2006; Michor et al., 2004). The immune system is adaptive and can therefore repeatedly attack new antigens even if old targets are eliminated. A more recent analysis investigated the systems-level cancer-immune interaction in a deterministic setting without taking into account adaptive evasion (Sontag, 2017). We previously considered an extreme version of this problem, namely an adaptive immune analogue capable of being permanently defeated by a single and durable stochastic evasion event (George and Levine, 2018). Here we consider a more realistic scenario and use it to interpret data regarding age incidence curves as well as tumor clonal evolution.

Specifically, we introduce a tractable mathematical framework for modeling the interaction between an adaptive T-cell repertoire capable of repeated recognition and a cancer population which may repeatedly evolve random mechanisms of immune evasion. We solve for the analytical escape and elimination probabilities and provide a statistical framework to study the mean-variance profiles of observed co-evolutionary recognition cycles. We apply our model to evolutionary data and predict ‘common initiation and rare progression,’ wherein a majority of cancer initiating events are controlled by an intact immune system, of which a small subset randomly escape.

## 2. Results

The following sections present the main findings of our analysis. Full mathematical details are provided in the SI.

### 2.1. Model Development

Our basic tumor-immune model consists of clones of cancer cells with pure birth at rate *r* until such time as they are detected by the immune system. Detection can occur once the tumor has reached a minimal detection size *m*_0_ and we work assuming either all clones are detected at size *m*_0_ with a one-time detection probability *q* (referred to as deterministic detection) or an ongoing chance of detection at sizes above *m*_0_ (adaptive detection). Once detection occurs, cells are killed at rate *d > r*, giving rise to a finite lifetime for that clone. There is also a probability per birth event *µ* ≪ 1 that the clone will give rise to an evasive tumor subclone. We previously discussed a model of this that assumed evasion events to be complete and durable against all current and future T-cells. Effectively, evading clones were rare but catastrophic cells which were invisible to the immune system, modeling an extreme event such as MHC-I down-regulation. In general, it is more reasonable to assume that most evasion events impart transient escape from recognition by the current dominant effector T-cell clone, with future detection possible by other clones.

Consequently, we study here the clonal evolution that results from repeated recognition and evasion events, referred to henceforth as *recognition cycles*. To do this, we assume that any evasive clone has in turn a chance of being detected when it grows to the detection size. A typical trace of the population dynamics generated by our model is shown in Fig. 1 where clones are created by evasion and die by detection over repeated cycles. Note that when the tumor population reaches some higher threshold *M* the cancer becomes clinically observable and “escapes” immune surveillance; in this simulation this occurs at around *t* = 50. Here the color code reflects subclones created during a specific cycle of the process. In general a given clone can give rise to multiple evasive lineages during its lifetime. For the deterministic detection scheme it is simple to argue (George and Levine, 2018) that the number of such daughter clones can be approximated by a Poisson distribution with parameter

**Figure 1:**
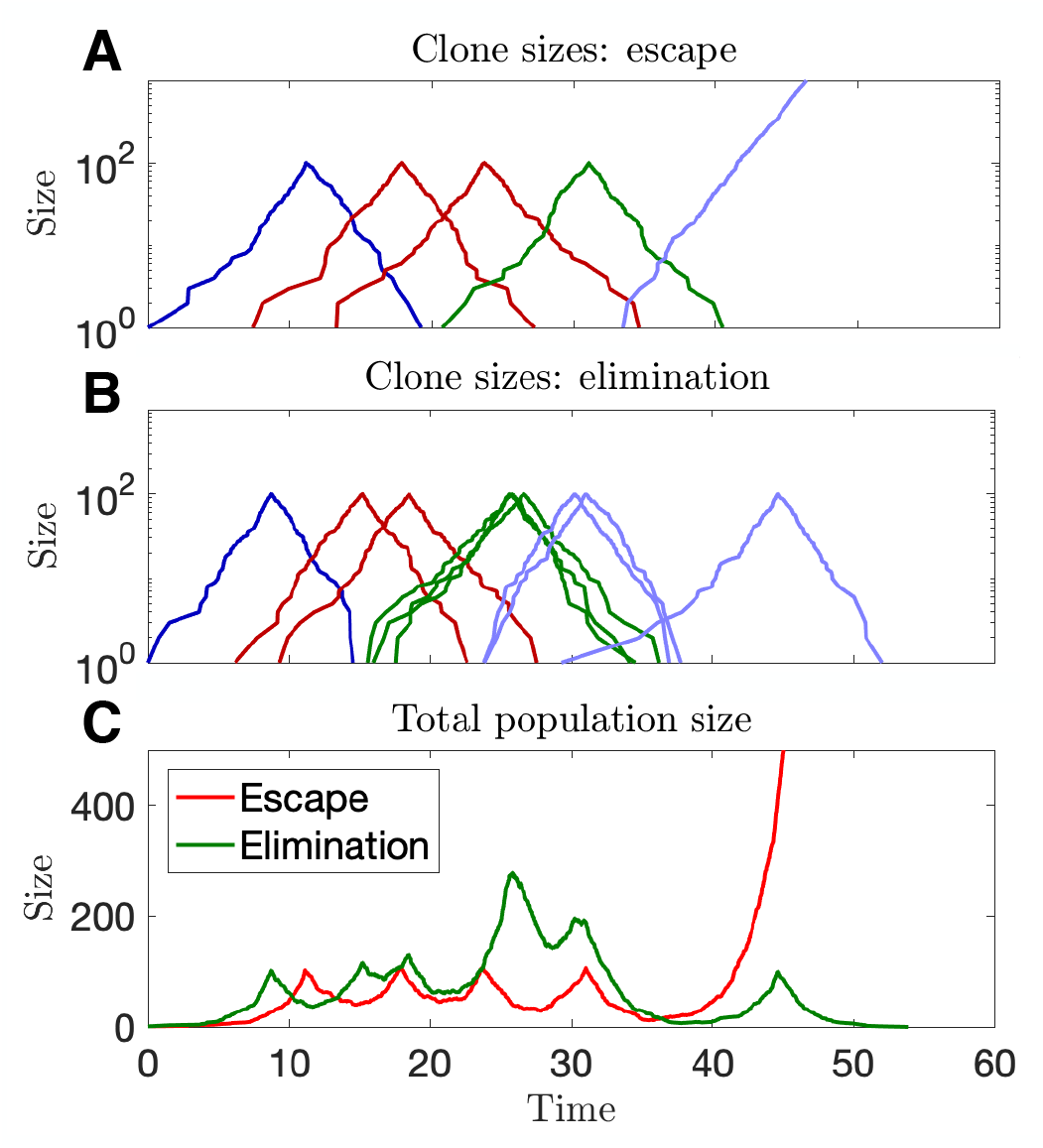
Overview of co-evolutionary trajectories. Simulations of the tumor-immune interactions depict clonal population sizes across recognition cycles (distinguished be different adjacent colors) in the event of ultimate escape (A) and elimination (B). The total population sizes are also recorded (C).

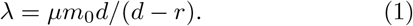

Thus the total evader “intensity”, i.e. the parameter governing the total number of clones produced at period *n*, is simply *γ*_*n,j*_ = *jλ* where *j* is the number of evasive clones at period *n*. A generalization of this formula for the adaptive assumption is presented in the SI. The model can therefore be framed as generalization of the Galton-Watson branching process, well-established in modeling biological systems (Athreya, 2006), with the addition of possible cancer escape. The process thus eventually culminates in either escape to size *M* or complete eradication of the tumor. We shall refer to the expected per-capita progeny as the branching parameter. In the deterministic case, this value is simply *λ* from Eq. 1.

Instead of tracking the population size at a given time, *t*, we focus instead on the number of evading clones, *Z*_*n*_ per recognition cycle, *n*, starting from one initial clone (Fig. 2A). The probability that a clone is recognized at cycle *n* is denoted by *q*_*n*_. This framework can in general handle any reasonable assumptions on the dependence of detection on cycle number. For example, it may be the case that the immune system becomes less capable over time due to having to deal with an increasing number of different clones. One could also generalize the above treatment to allow for the recognition rate to depend explicitly on the number of clones present at cycle *n* or even on the individual evasive clone as a function of its appearance order. For simplicity, we will mostly assume that *q*_*n*_ is a constant which will be denoted as *q*_*c*_. These assumptions lead to a Markov process relating the number of clones at cycle *n* to the number of clones at cycle *n* + 1 (Fig. 2B,C). The probability of generating *k* clones in the next cycle given the existence of *j* clones in the current cycle is

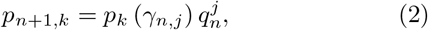

with

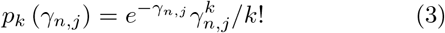

**Figure 2:**
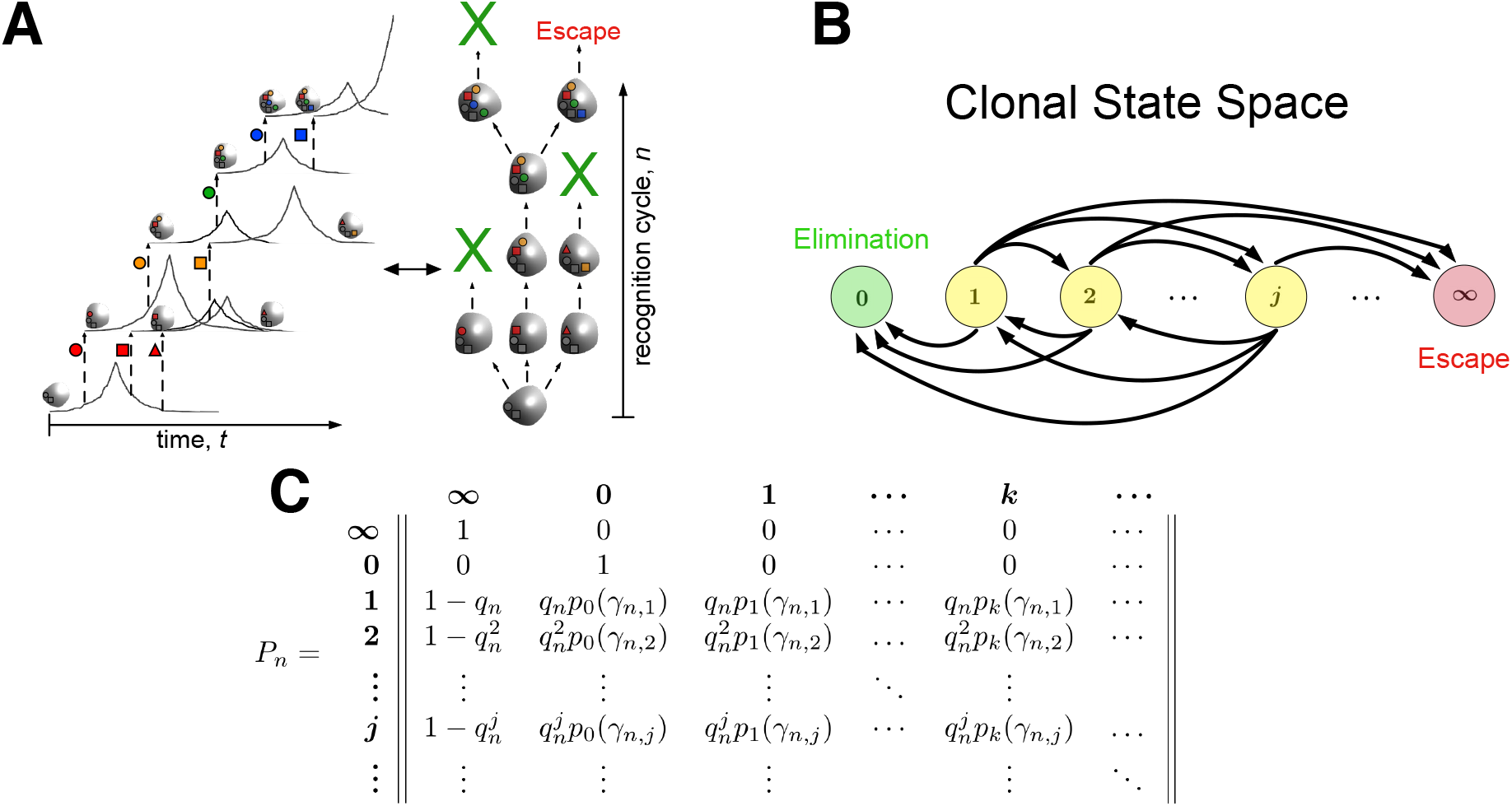
Cancer-immune co-evolutionary dynamics. (A) A single initial clone grows, becomes recognized, and is eliminated, but not before producing immune evasive clones (distinguished by shapes), which impart transient immunity against the current effector T-cell clone. We assume that each evader has an independent opportunity to either escape or become recognized by the T-cell repertoire and may also give rise to additional evaders (evasion events occurring within the same recognition cycle share the same color and are differentiated by shape). (B) The state space of this discrete process is the number of distinct evasive clones at each generational period. Escape (state ∞) and elimination (state 0) are absorbing states, and all other intermediate states communicate; (C) This model may be represented by an inhomogeneous Markov process with a transition probability matrix that depends on the immune clearance probability of each of the *j* clone at time *n*, 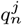, and the probability of generating *k* clones from *j* current clones, *p*_*k*_(*γ*_*n,j*_) given by Eq. 3.

The immune system must recognize every subclone at each period or else the threat escapes. Thus, for *j* clones at period *n*, the escape probability is

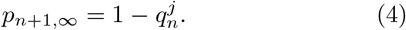

Finally, the probability of tumor elimination is just the probability that no evasive clones are generated and no escape occurs, namely

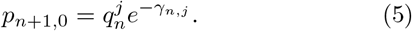

### 2.2. Escape and elimination probabilities

Let *E*_*n*_ (resp. *F*_*n*_) be the event that the threat escapes (resp. is eliminated) at period *n*. One can directly derive a backwards discrete-time master equation that determines the long-time limit of these splitting probabilities. Specifically focusing on the elimination and taking *q* = *q*_*c*_ to be constant, we have

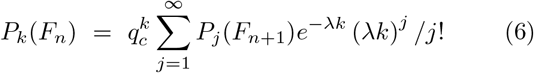

where *P*_*k*_(*F*_*n*_) is the probability that a tumor trajectory having *k* clones at cycle *n* eventually leads to complete tumor elimination. Clearly each clone is independent from its future trajectory and hence *P*_*k*_(*F*_*n*_) = *P*_*1*_(*F*_*n*_)^*k*^. Thus Eq. 6 becomes a closed-form equation for *P*_*1*_(*F*_*n*_)

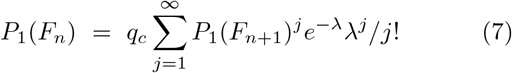

Taking the steady-state limit of this equation yields an equation for the asymptotic splitting probability *p*^∗^

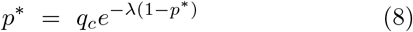

One can similarly find the asymptotic value of the escape probability and verify that it is just 1 −*p*^∗^; in other words the system always chooses one of the absorbing states in the long-time limit. A more complete treatment presented in the SI allows this result to be extended to evaluate the full time dependent value of *P*_1_(*F*_*n*_) and also consider more general assumptions for the *q*_*n*_.

The above outlines the model for the deterministic case since detection, should it occur, must do so at size *m*_0_. Adaptive detection incorporates immune decline via an immunomodulatory parameter *ν >* 0 that results in larger average detection sizes (See SI for full details). This analytical framework above agrees well with results obtained from simulating the full co-evolutionary process (Fig. 3A,B). ‘Escape by underwhelming’ is an important experimentally observed feature wherein threats of intermediate growth rates are at a survival disadvantage relative to their faster and slower growing counterparts (Bocharov et al., 2004). Our model recapitulates this behavior for minimal detection thresholds *m*_0_ limited by the total cancer growth rate (Fig. 3C), an assumption known as the ‘growth-threshold conjecture’ (Arias et al., 2015). This simple theoretical framework generates a significant degree of diversity in dynamical behavior, governed completely by the clearance probability and the branching pa-150 rameter (Fig. 3D).

**Figure 3:**
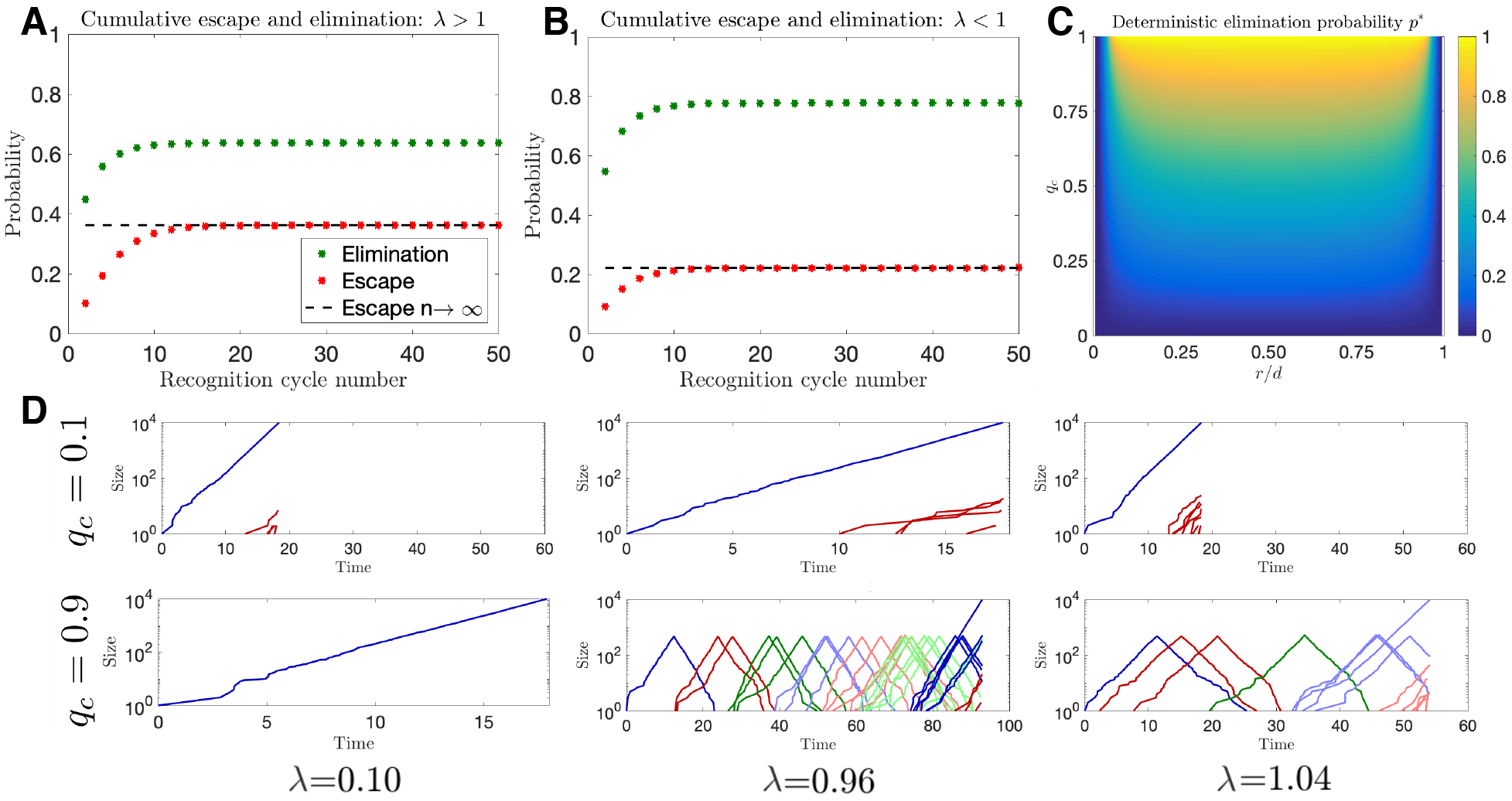
General dynamics of the tumor-immune co-evolutionary model. Escape and elimination probabilities are plotted as a function of recognition cycle number for (A) supercritical (*λ* = 1.04) and (B) subcritical (*λ* = 0.96) branching (in each case, *q*_*c*_ = 0.95 and simulations are averaged over 10^5^ iterations); (C) The dynamics induced by Eq. 8 permit a unique limiting elimination probability, *p*^∗^, which is plotted as a function of clearance probability and relative net growth rate (*λ* calculated using Eq. 1 with *µ* = 10^*−*6^, *d* = 0.2, *R* = 10^4^, and *m*_0_ = *R/r*); (D) A variety of evolutionary trajectories are plotted conditional on ultimate escape, assuming various clearance and branching parameters (in all cases, deterministic recognition was assumed with *m*_0_ = 500 *r/d* = 0.5).

### 2.3. Differences in cancer early age incidence correlate with evasion rate

As proof-of-principle that local physiological changes in immune function may affect observed cancer frequency, even among healthy individuals, we study the effects of physiological declines in immune turnover on increases in early age incidence for various cancer types, using large population publicly-available datasets (Fig. S6) (Cancer Research UK). Previous studies have shown that age-incidence data can be fit to simple models assuming a declining immune system with age (Palmer et al., 2018). Our dynamical model predicts, all else being equal, that tumor escape likelihoods vary directly as a function of evasion rates *µ* implicitly through the branching parameter, and although evasion need not be limited exclusively to genomic mechanisms, we use experimentally derived per-cell mutation rates obtained from a large pan-cancer analysis (Lawrence et al., 2013).

We calculate the slope parameter of linear regression for early age incidence (Figs. 4A, S6) and find a strong correlation between these values and the evasion rate. Aggregate breast cancer appears to be an exception, perhaps a result of early screening detection of hormone-sensitive disease. Restricting the comparison to triple negative breast cancer gives data that is more consistent with the overall the trend (Fig. 4B). Decreases in immune function modeled via an increased adaptive parameter *ν* (see SI), assumed directly proportional to early age increases, result in increased ultimate escape probability, 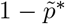. We find excellent agreement in the predicted incidence slope-*µ* profile for the best-fit scaled range of immune function parameter *ν* (Fig. 4C). Moreover, using this normalized range for *ν* in the model predicts a linear increase in incidence for all observed mutational rates (Fig. 4D). Together, these findings demonstrate that cancer progression and escape are well explained by variance in immune function across nearly all cancer types.

**Figure 4:**
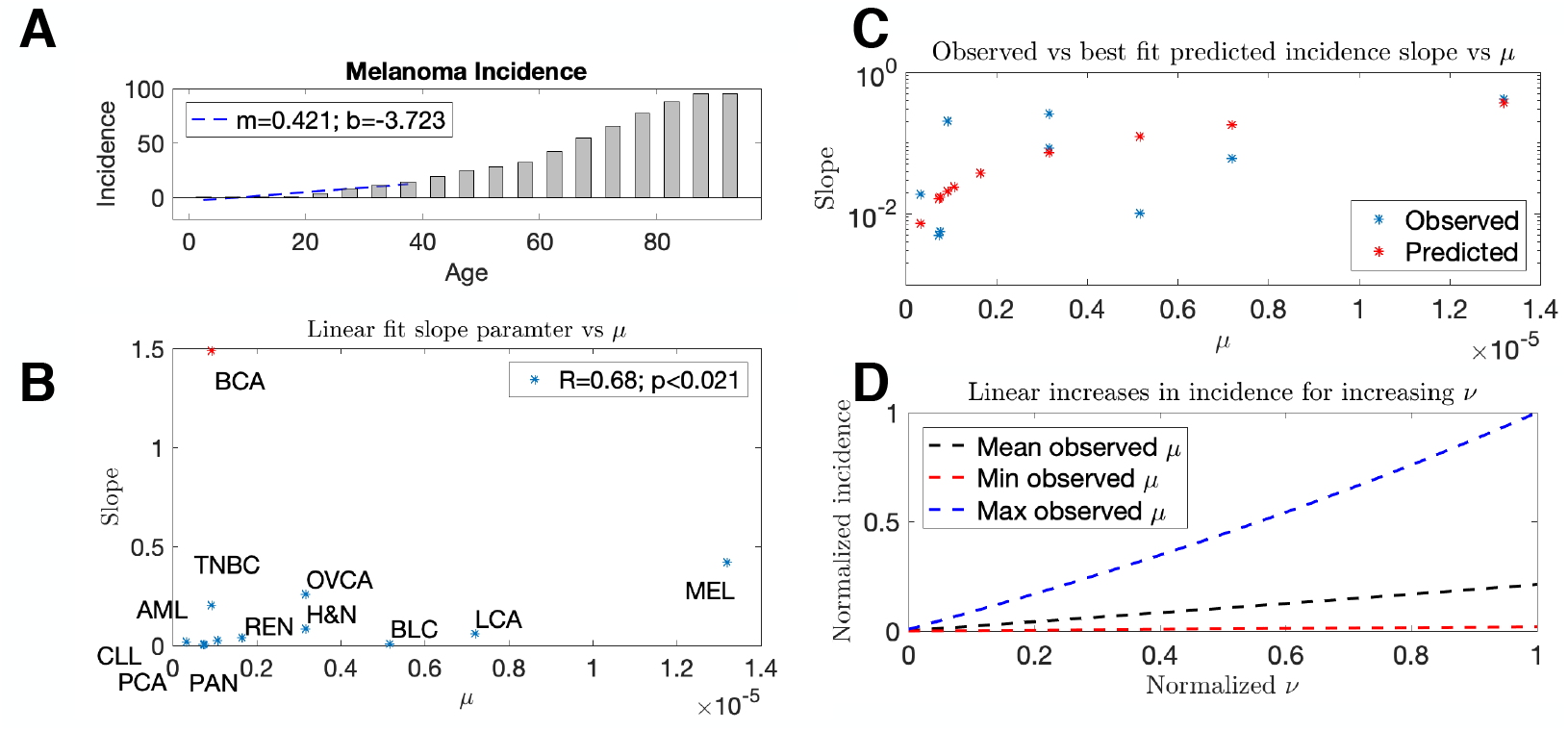
(A) The least-squares linear regression slope parameter of early age cancer incidence (between ages 0 and 40 years) is calculated for a variety of cancer subtypes (melanoma shown); (B) Rate of change in cancer early age incidence vs. evasion rate. For each age incidence curve, linear regression is performed for incidence between ages 0 and 40 years. This parameter is plotted as a function of per-cell mutation rates for each cancer type; (C) Assuming that the relationship between age and *ν* is linear, age=*kν*, the slope in cancer age-incidence can be calculated as a function of increasing *ν*, and are plotted for *k* that gives the least-squares error as compared to the experimental data; (D) This parametrization consequently predicts linear increases in early cancer incidence as a function of diminished immune performance for all observed ranges of evasion rates *µ*.

### 2.4. Evolutionary timing of renal clear cell carcinoma predicts prolonged co-evolution with common tumor incidence and rare progression

Given the ability of our mathematical model to link observed cancer evasion and recognition cycle timing to underlying tumor-immune co-evolutionary dynamics, we next assessed the extent of immune evasion and clearance observed in cancer evolutionary data. The TRACERx renal clear cell carcinoma (RCCC) dataset (Mitchell et al., 2018) was particularly useful, providing estimates of the timing of landmark evolutionary events. Surprisingly, and consistent with model behavior under sustained control (Fig. 3D), the authors found on average extended periods of clonal evolution sustained by small population sizes (estimated to be of order 10^2^ cells). Using the time of most recent common ancestor (MRCA) arrival as a representative of escape, we estimated the expected number of recognition cycles given the observed intervening time from MRCA detection time and from disease initiation vs MRCA. If *t*_*m*_ is the time it takes a clone to grow to minimal detection size *X*(*t*_*m*_) ~ 10^2^ and *T*_*M*_ the times it takes to growth to ultimate detection size *X*(*t*_*M*_) ~ 10^9^, then the intervening time assuming *n* recognition cycles occurs may be estimated as ∆*t* = *t*_*M*_− *nt*_*m*_. From this, we can estimate the number of recognition cycles via

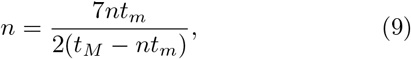

where *nt*_*m*_ and *T*_*M*_ are the observed times (see SI for details). Our calculations predict that, on average, RCCC undergoes approximately 27 distinct, immunologically relevant recognition cycles prior to escape (Fig. 5A-B).

The parameter range that generates this behavior is near criticality (*λ* ≲ 1) with high clearance probabilities (*q*_*c*_ approaching 1), suggesting effective immune handling of renal cell threats (Fig. 5C). To further investigate the extent of immune protection, we used parameter estimates consistent with these findings to assess the frequency of observed cancer incidence relative to the total number of (unobserved) cancer initiating events, and predict that initiation is 18 times as frequent as measured incidence. Together, these results suggest that the adaptive immune system actively filters many potential threats manifesting as *de novo* initiating cancer clones, and may occasionally miss due to statistical chance, in support of the ‘bad-luck hypothesis’ (Tomasetti and Vogelstein, 2015). The resulting population dynamics closely resemble those observed in the cancer incidence data, with a large disease period consisting of clonal evolution with low tumor burden, followed by either elimination or escape with extensive sub-clonal evolution (Fig. 5D).

**Figure 5:**
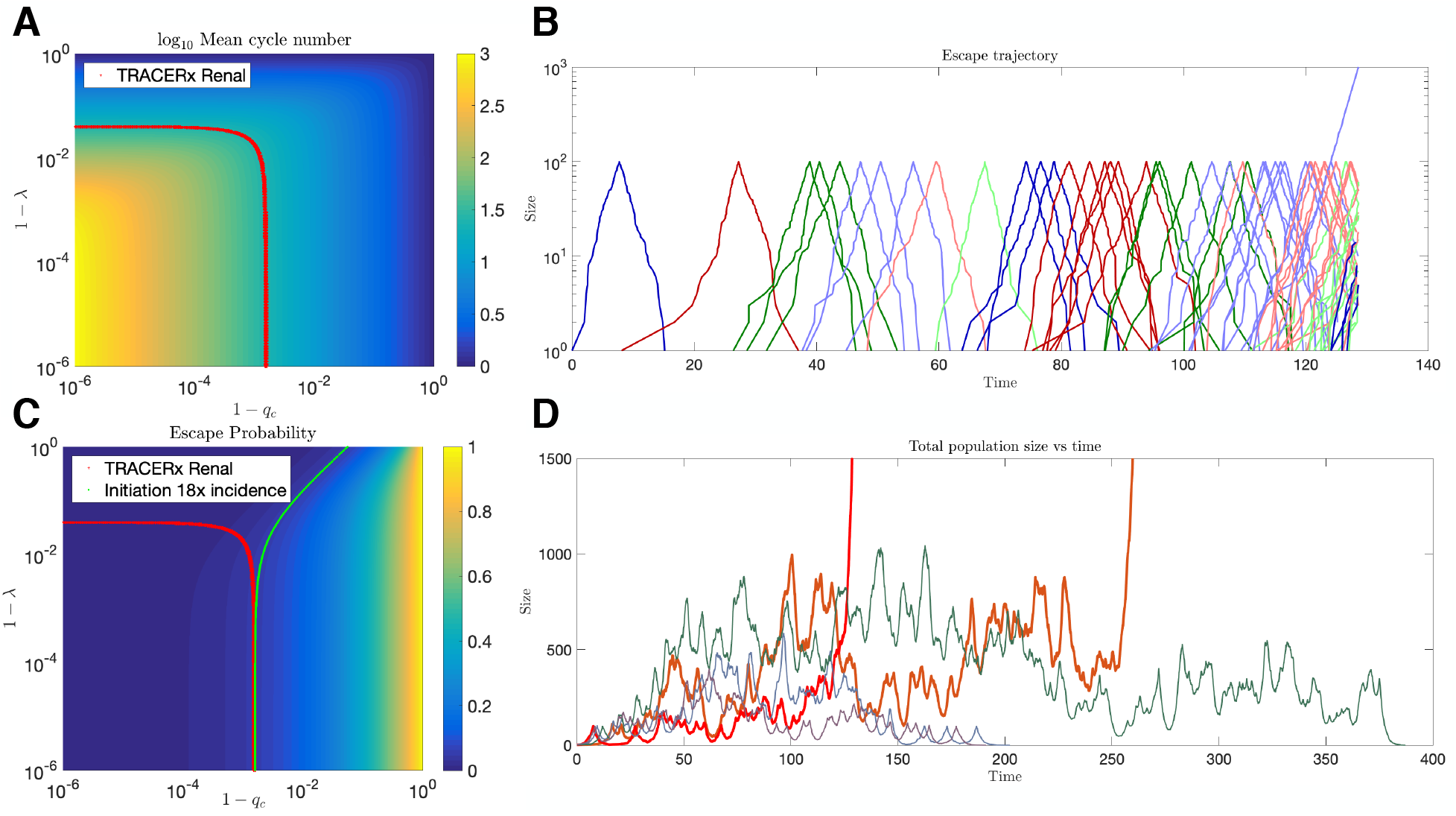
Evolutionary dynamics of renal clear cell carcinoma. (A) Relative timing of the TRACERx Renal time to MRCA vs time to clinical escape implies an average of 26.6 cycles numbers, which restricts the range of allowable values of *q*_*c*_ and *λ* in the co-evolution model (red line traces the contour line for relevant mean cycle number); (B) A typical stochastic realization of the clonal branching that occurs in the deterministic case before and after escape (colors distinguish adjacent subclones arising within the same cycle number); (C) minimal values of observed clearance probability provide an upper estimate for the observed escape probability at 1/18 (plotted as a function of *q*_*c*_ and *λ*; green line traces the contour line for relevant escape probabilities); (D) Simulation output of the total population size as a function of time for various realizations of the process.

### 2.5. Fluctuations in lung cancer preceding escape partition disease subtypes based on immune function

Given the predicted implications of immune perfor-mance on eliminating early and indolent disease, we wanted to investigate the relationship between disease subtype and predicted immune status. Available data on large cohort lung cancer (LCA) patients (Jamal-Hanjani et al., 2017) provided enough samples for each studied subtype to estimate mean and variance profiles for the distribution of clonal mutational events. This analysis was repeated for each LCA subtype and then compared to all predicted values in the model. Since only a small (unknown) fraction *α* ≪ 1 of total mutations are expected to impart immune evasion, we consider the resulting allowable mean-variance frontier assuming a range of *α* small. Very briefly, for *X* being the total number of clonal non-synonymous mutational events with respective mean and variance given by *µ*_*X*_ and 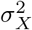, the mean and variance for the fraction *αX* which are relevant to immune evasion relate to the observed parameters *µ*_*X*_ and 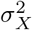 via

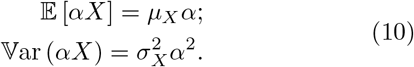

Eqs. 10 define a quadratic mean-variance curve parameterized by *α* which can be applied to compare the observed curves to all allowable simulated parameters in the domain considered in Fig. 5A,C to determine model conditions which are consistent with empirical data. Our above model defines a baseline relationship that well-characterizes the reported mean-variance profiles across all LCA evolutionary data subtypes (Fig. 6). However, several variations in the mean-variance frontier are only achievable in our generalized model assuming either immune suppression or heightened immune surveillance (See SI for full details). Interestingly, squamous cell carcinoma, and to a lesser extent positive smoking status, are more consistent with a process which is significantly immunosuppressed, suggesting that the observed pattern of mutational variability emerges in the setting of reduced immune targeting efficiency. Intriguingly, the predicted mean-variance frontier requires enhancements to immune clearance rates over successive periods in order to explain the non-smoker signature (Eq. S123).

**Figure 6:**
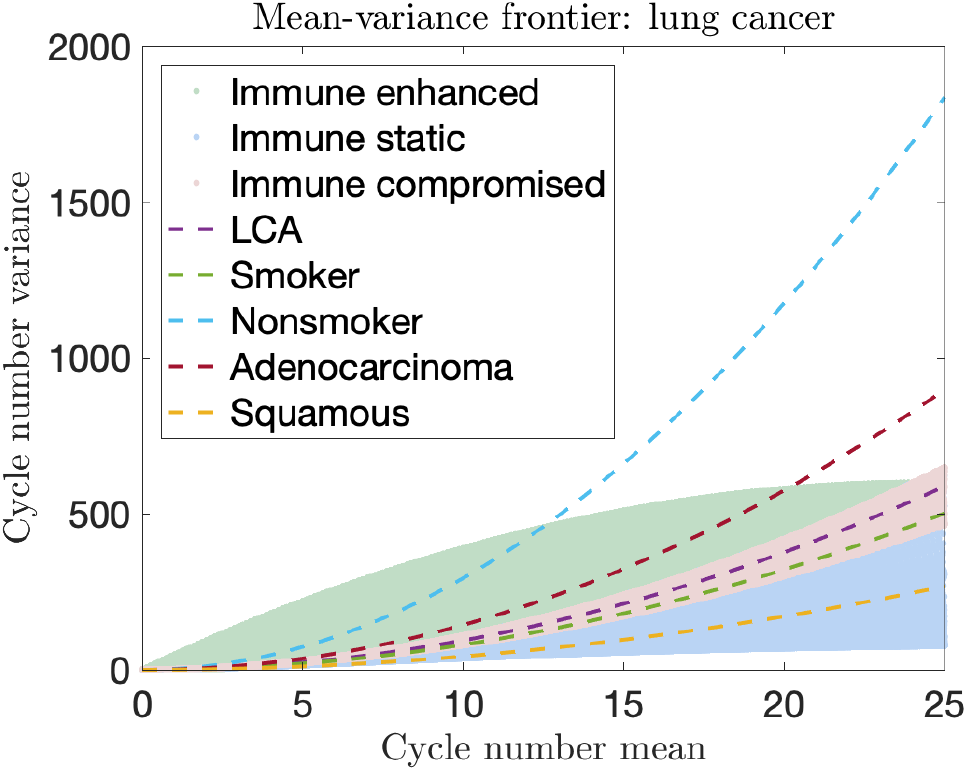
Mean-variance frontiers for lung cancer subtypes. Empirical mean-variance frontiers are plotted using Eq. 10 and observed data for various LCA subtypes (dashed lines). These are compared to predicted allowable regions in the mean-variance space based on all parameter combinations studied in Fig. 5A,C (colored regions). Red region corresponds with allowable frontiers assuming deterministic recognition (*ν* = 0). Blue region highlights areas of non-overlap in adaptive recognition (*ν* = 10 taken to be 10% of *m*_0_). Green regions represent dynamics assuming immune enhancement over time (*p*_*c,n*+1_ = max{*p*_*c*,∞_, *p*_*c,n*_ + (*p*_*c,n*_*p*_*n*,∞_/25)} for *p*_*c*,∞_ = 0.9).

## 3. Discussion

Understanding the underlying early clonal structure of an evolving threat under adaptive immunosurveillance is critically important for better predicting the extent of tumor immunoediting, estimating the frequency of disease progression and hence immune targeting efficacy, and proposing optimized treatment strategies conditioned on clonal distributions following immune escape. Here, we developed a stochastic population dynamical model of the battle between an evolving cancer population and the CD8+ T-cell immune repertoire. The underlying dynamical behavior of the model subsequently encodes information on both the likelihood and timing of ultimate cancer escape or elimination. For reasonable immune parameter estimates, our model predicts that the disease state ultimately results in either escape or elimination with certainty, given sufficient time. We use our mathematical framework to calculate the likelihood of these events as function of cancer growth and evasion rates, and immune killing and detection rates. We further demonstrate that incorporation of the growth-threshold conjecture results in empirically observed ‘escape by underwhelming.’ Our framework is largely consistent with the importance of immunosurveillance in early disease progression as evidenced by differential increases in cancer incidence that scale with evasion rate across nearly all cancer types.

Using the excellent data available on the timing of early evolutionary events in RCCC, together with the observation that early clonal evolution is estimated to be sustained at small cell sizes, we show that the mean number of recognition cycles is quite large, suggestive of a drawn-out competition between immune and cancer compartments. Our results suggest that cancer progression from early initiation to appreciable disease only occurs a small fraction of the time, in support of the bad-luck hypothesis wherein a majority of observed cancer cases in healthy individuals is due to random chance. We remark that these results are most sensitive to the assumed minimal population size (here taken to be *m*_0_ = 10^2^ as estimated empirically), the resulting mean cycle number estimate scales as a decreasing function of *m*_0_ as 2/log_10_*m*_0_. Despite this, significant and prolonged co-evolution is still predicted (15 recognition cycles in renal cancer patients) assuming an order of magnitude increase in the minimal detection size. Though empirically challenging, further investigation into the early tumor-immune interaction following initiation could refine this estimate and shed light on the underlying evolution of early disease. When comparing the mean-variance frontiers in the clonal distributions of TRACERx LCA subtypes, we found that squamous cell carcinoma and positive smoking status were consistent with simulated profiles assuming immune impairment, while non-smokers and adenocarcinoma followed an opposite trend. Our results demonstrate the value of further experimental investigation into the benefit of distinct treatment strategies based on cancer subtype and immune function of each patient.

Although we have focused our analysis on describing the control and progression of cancer, our model is broadly applicable in understanding similar phenomena in other intracellular infection which evolve mechanisms of immune evasion. In HIV, for example, the delayed minimal disease burden produced by repeated recognition cycles in our model shares some general features of the latent period. Future efforts to apply this model in a broader context may provide insights to the observed dynamics of slow-progression intracellular threats.

## Author’s Contribution

JTG conceived of and designed the research, analyzed the data, contributed new analytic equations, and wrote the manuscript. HL designed research, analyzed the data, contributed new analytic equations, and wrote the manuscript. Both authors have approved the final article.

## Acknowledgments

We would like to thank Jeffrey Molldrem for fruitful discussions on immunotherapy and cancer-immune coevolution. JTG is supported by the National Cancer Institute of the National Institutes of Health (F30CA213878). HL is supported by Physics Frontiers Center NSF Grant PHY-1427654 and by NSF PHY-1605817.

## Appendix

Supplementary information with full mathematical details are provided in the attached document.

